# Preliminary First-in-Human Pharmacokinetic Evaluation, Dosimetry and Safety of 7-[^18^F]-Fluorotryptophan as PET Imaging Agent to Visualize Serotonin Synthesis in the Brain

**DOI:** 10.1101/2025.08.26.672460

**Authors:** Sophie Stotz, Rodrigo Moraga-Amaro, Boris D. Zlatopolskiy, Michael Gemki, Gitte M. Knudsen, Bernd Neumaier, Cristian Soza-Ried, Horacio Amaral, Vasko Kramer, Matthias M. Herth

## Abstract

Tryptophan (Trp) is the precursor for serotonin synthesis and other biologically relevant metabolites. We evaluated the novel radiotracer 7-[^18^F]Fluorotryptophan (7-[^18^F]FTrp) to assess its biodistribution, dosimetry, and potential for imaging brain Trp metabolism in humans. Six healthy volunteers underwent whole-body PET/CT imaging over 5.5 hours following intravenous injection of 7-[^18^F]FTrp. An additional four subjects underwent dynamic brain PET imaging for 2 hours. Time-activity curves (TACs) were extracted for source organs using VOIs defined on co-registered CT and PET images, and dosimetry was calculated using OLINDA software. The radiotracer showed rapid uptake and distribution, with highest activity observed in the liver, pancreas, salivary glands, in combination with urinary excretion. Brain pharmacokinetic analyses with image-derived input function (IDIF) determined that Patlak analyses were the best fit for brain image analyses. Brain uptake was modest, with highest region-specific accumulation in the pineal gland, which is a known site for serotonin synthesis. The estimated effective dose was within the expected range for ^18^F-labeled compounds (14.1 ± 0.2 μSv/MBq). Our findings indicate that 7-[^18^F]FTrp is safe for human use, demonstrates favorable kinetics for studying both brain and peripheral Trp metabolism, and warrants further exploration in patients with serotonin metabolism disorders.

## Introduction

Serotonin (5-hydroxytryptamine, 5-HT) is a key neurotransmitter that plays a critical role in regulating mood, cognition, and a variety of physiological processes, including sleep, appetite, and gastrointestinal function. Alterations in serotoninergic neurotransmission have been linked to several neuropsychiatric and neurological disorders, such as depression, anxiety, schizophrenia, and Parkinson’s disease, highlighting the importance of studying serotonin pathways in the human brain.^1,2^ Serotonin synthesis is initiated from the essential amino acid tryptophan (Trp) in both the gut and the brain, which is converted into 5-hydroxytryptophan by the enzyme tryptophan hydroxylase (TPH), and subsequently into serotonin by aromatic L-amino acid decarboxylase.^3^ The neuronal isoform of TPH, TPH2, is mainly expressed within the raphe nuclei, while in the periphery TPH1 is expressed the pineal gland and key endocrine organs like pancreas and the gastrointestinal tract.^4-8^ Alternatively to the serotonin pathway, Trp enters the kynurenine pathway which is highly upregulated under inflammatory conditions.^9^ Given the fundamental role of Trp in serotonin production, variations in tryptophan availability and metabolism can significantly influence serotoninergic function. Inadequate levels of Trp, like serotonin levels, have been associated with various neuropsychiatric disorders, including depression, schizophrenia, and Parkinson’s Disease.^10^ Trp supplementation has been explored as a therapeutic strategy for boosting serotonin levels in conditions like depression and sleep disorders.^11^ As such, the involvement of Trp in serotonin synthesis and its dysregulation in neurological conditions requires understanding its biodistribution, metabolism, and dynamics in the brain.

In consequence, the clinical need for reliable imaging techniques to assess serotonin synthesis and metabolism has grown due to its potential as a predictive biomarker for serotonin dysregulation.^12^ Positron emission tomography (PET) is as a valuable tool for non-invasive imaging of neurotransmitter systems, allowing for the assessment of serotonin synthesis, receptor binding, and transporter function in the living brain.^13^ However, the development of radiotracers that accurately reflect serotonin synthesis and turnover has proven challenging. Existing radiotracers such as α-[^11^C]methyl-L-tryptophan have been employed to visualize serotonin synthesis, but they suffer from limitations such as non-specificity, slow brain uptake, and complex kinetic modeling.^14,15^

7-[^18^F]Fluorotryptophane (7-[^18^F]FTrp), a novel radiolabeled Trp derivative, represents a promising advancement in serotonin synthesis imaging.^16^ Fluorine-18 (^18^F) offers superior imaging properties compared to carbon-11, including a longer half-life (110 minutes versus 20 minutes) and higher spatial resolution, making it suitable for widespread clinical applications. While other fluorinated Trp derivatives undergo rapid defluorination in vivo, 7-[^18^F]FTrp is stable and has proven promising uptake in the pineal gland and the raphe nuclei in rats.^16^ Preclinical studies of 7-[^18^F]FTrp have demonstrated favorable biodistribution, brain penetration, and accurate delineation of kynurenine and serotonin-producing organs from background tissues, indicating its potential as a radiotracer for serotonin synthesis imaging.^17^ Furthermore, the ^18^F label enables more extended imaging sessions, allowing for dynamic studies of serotonin metabolism over time.

In this study, we report on the dosimetry and biodistribution of 7-[18F]FTrp, along with the kinetic analysis of the first dynamic brain PET imaging study in humans. This work builds upon preclinical evidence by translating the tracer into a clinical setting, providing initial data on its safety, pharmacokinetics, and potential as a tool for serotonin imaging in neuropsychiatric and neurological disorders.

## Materials and methods

### Radiochemistry

7-[^18^F]FTrp was synthesized under Good Manufacturing Practice (GMP) compliant conditions and according to published protocols.^16^ In brief, 7-[^18^F]FTrp was produced using an automated procedure and standard IFPs for nucleophilic labeling (Synther+, IBA Radiopharma Solutions, Louvain-la-Neuve, Belgium). 40-60 GBq of F-18 fluoride (IBA cyclone 18/18) was trapped on a strong anion exchange cartridge (QMA Light, Waters Accel Plus QMA Light cartridge, preconditioned with 10 mL 0.5M potassium carbonate, 10 mL water, and 3 x 10 mL air) and eluted with eluent from vial 1 (1.5 mg tetraethyl ammonium bicarbonate in 1.0 mL methanol) into the reactor. The fluoride was dried by azeotropic distillation, the precursor (5.0 mg Boc-7-BpinTrp-OtBu, 7.0 mg tetrakis(pyridine)copper(II) triflate, 0.6 mL dimethylacetamide anhydrous, 0.25 mL n-butanol) was added from vial 2, and the reaction mixture was heated to 110 °C for 10 min. The reaction was quenched with 2.5 mL water, transferred to a second IFP, trapped on a C18 cartridge (C18 light short, preconditioned with 10 mL ethanol, 10 mL water, and 3 x 10 mL air) and eluted into reactor 2 with 1.5 mL acetonitrile from vial 1. Concentrated hydrochloric acid (1.0 mL; 30%) was added from vial 2 and the resulting mixture was heated to 60 °C for 5 min, diluted with 2.5 mL water from Vial 4, and transferred to the HPLC loop via an airlock filter (Millex Nylon NH, 0.45 μm, Merck Millipore). The product was then purified via semi-preparative HPLC (Phenomenex, Luna 5 μm C18(2) 100 Å, 250×10 mm; mobile phase: 8% ethanol in 30 mM sodium acetate buffer, pH 4.3) at a flowrate of 4 mL/min. 7-[18F]FTrp was collected after 20 min into a vial containing 18 mL 0.9% sodium chloride and the final solution was passed through a sterile filter and obtained as a solution ready for injection. The radiochemical and chemical purity and identity was tested by radio-HPLC and radio-TLC and further quality control was performed (radionuclide purity, residual solvents, appearance, pH, endotoxins, sterility, filter integrity) in compliance with current pharmacopeias. 7-[^18^F]FTrp was obtained in radiochemical yields of 5-10 % after a total synthesis time of 105-115 min and the radiochemical purity was ≥ 95 %.

### Subjects

The dosimetry study was performed in accordance with the regional ethics committee (CEC SSM Oriente, permit 20140520) and written consent was received from all participants. Six healthy controls were included and participated in the dosimetry protocol, comprised of 3 females and 3 males, with a mean age and body mass index, respectively, of 36 ± 6 years and 28.7 ± 2.7. For the dynamic brain study, 4 healthy controls were included (2 females and 2 males) with a mean age of 38 ± 10 years and mean body mass index of 25.5 ± 1.5. Detailed demographics are shown in Table 1. All subjects were healthy and free of any medical condition at the time of the scans (determined by medical examination and questionnaire) and fasted on the day of the study.

**Table 1:** Demographics of the subjects that underwent a dosimetry scan (Dos) and the dynamic brain study (Brain).

| Subject ID | Sex | Age (y) | Height (m) | Weight (kg) | BMI (kg/m<sup>2</sup>) | Injected activity (MBq) |
| --- | --- | --- | --- | --- | --- | --- |
| DOS_001 | M | 38 | 1.77 | 75 | 23.9 | 223.5 |
| DOS_002 | M | 44 | 1.82 | 99 | 29.9 | 229.0 |
| DOS_003 | F | 40 | 1.60 | 80 | 31.2 | 236.1 |
| DOS_004 | F | 27 | 1.67 | 84 | 30.1 | 244.2 |
| DOS_005 | M | 33 | 1.70 | 76 | 26.3 | 212.4 |
| DOS_006 | F | 34 | 1.63 | 82 | 30.9 | 205.7 |
| **Mean ± SD** |  | **36 ± 6** | **1.70 ± 0.08** | **82.7 ± 8.0** | **28.7 ± 2.7** | **225.2 ± 14.4** |
| BRAIN_007 | M | 42 | 1.68 | 72 | 25.5 | 226.1 |
| BRAIN_008 | M | 50 | 1.87 | 88 | 25.2 | 232.0 |
| BRAIN_009 | F | 27 | 1.64 | 74 | 27.5 | 226.4 |
| BRAIN_010 | F | 34 | 1.50 | 54 | 24.0 | 207.2 |
| **Mean ± SD** |  | **38 ± 10** | **1.67 ± 0.15** | **72 ± 14** | **25.5 ± 1.5** | **222.9 ± 10.8** |

### PET/CT and PET/MR imaging

210.0 ± 41.4 MBq (dosimetry study) and 222.9 ± 10.8 MBq (brain study) of [^18^F]FTrp was administered intravenously to the subjects.

Quality control of the PET/CT scanner was performed daily before every study. For the participants of the dosimetry study, a series of 14 whole-body (WB) PET/CT scans (Biograph Vision: n = 3 subjects, Biograph mCT Flow: n = 3 subjects, both Siemens, Erlangen, Germany) covering head to mid-thigh (10–11 bed positions) was acquired over the course of 5.5 h and in three separate sessions as shown in table 2. For the participants of the brain study, a dynamic PET/CT scan (Biograph Vision: n = 2 subjects, Biograph mCT Flow: n = 2 subjects) covering only the head was acquired over the course of 2 h.

**Table 2.** PET scanning protocol.

| *Session* | *Duration* | *No of PET scans* | *Bed position time* |
| --- | --- | --- | --- |
| 1 | 0 - 120 min | 10 | 4 x 30s<br>4 x 60 s<br>2 x 120 s |
| Break | 120 – 180 min | - |  |
| 2 | 180 – 225 min | 2 | 2 x 270 s |
| Break | 225 – 285 min | - |  |
| 3 | 285 – 330 min | 2 | 2 x 270 s |

Subjects were asked to void their bladder in the break after each imaging session. A low-dose CT scan was performed for attenuation correction and co-registration prior to each session. PET data were corrected for random coincidences, normalization, attenuation, scatter, and dead-time losses. The data were reconstructed using an ordinary Poisson ordered subset expectation maximization(OP-OSEM) 3D iterative algorithm (Vision: 9 iterations, 5 subsets, matrix: 220 × 220; mCT Flow: 2 iterations and 21 subsets, matrix: 200 × 200) applying time-of-flight (ToF) and point-spread function (PSF) modelling with post-reconstruction smoothing (Gaussian,4 mm full-width at half-maximum). The dynamic PET images were reconstructed using the same algorithm and user-defined dynamic framing (44 frames, 120 min: 6 x 10 s, 6 x 20 s, 7 x 60 s, 5 x 120 s, 20 x 300 s).

### PET image analysis

After reconstruction, PET images were motion-corrected in atlas space. PET images were analyzed using PMOD (version 3.9, Bruker, Karlsruhe, Germany). Voxels of Interest (VOIs) were manually drawn according to the CT images (brain, kidneys, lungs, liver, pancreas, spleen, stomach, left and right colon, skeleton and femora, total heart, red marrow in the lumbar vertebrae L2-L4) and adjusted using threshold-based segmentation if appropriate. The VOIs were transferred to the corresponding, co-registered PET images with minor manual adjustment if needed. For urinary bladder and salivary glands, VOIs were outlined by threshold-based segmentation on each individual PET scan. The remainder was obtained by subtracting the organ VOIs from a total body VOI based on each individual PET scan to control for movement in between PET scanning sessions. Not all VOIs were directly used as source organs for dosimetry calculations. Total activity in red marrow was calculated using the activity from red marrow VOIs outlined for L2-L4 vertebrae assuming that L2-L4 constitute 60% of total lumbar vertebrae volume and the lumbar vertebrae contain 12.3% of the total red marrow mass.^18^ For calculation of total bone activity, the femora were excluded from the skeleton VOI by subtraction. As the legs represent 33.6% of total bone mass, the obtained activity was scaled by a factor of 1.506 (1/0.664).^18,19^ The activity of the red marrow was subtracted. The total bone activity was then distributed between cortical bone mineral surface (75%) and trabecular bone mineral surface (25%).^19^ The activity in the intestine was approximated by adjusting the total activity according to the percentual organ weight provided by OLINDA (left/right intestine: total activity * 0.158, small intestine: total activity * 0.684). Finally, decay corrected time-activity-curves (TACs) were generated for all source organs and the total body VOI.

Dosimetry calculations were performed in OLINDA (Organ Level INternal Dose Assessment version 2.2) using the previously obtained TACs. All TACs were fitted depending on the degree of correlation to a mono-, bi- or tri-exponential function, excluding time points if necessary for the fit. After fitting all source organs, kinetics, organ doses, and effective dose were estimated automatically using an ICRP adult male and adult female model for male and female healthy controls, respectively, with the organ weight adjusted to the actual weight of the participants.

### PET brain pharmacokinetic analysis

After image acquisition in LIST-mode, the raw data was iteratively reconstructed as described above, corrected for attenuation, scatter and decay of radioactivity. For data analysis, each PET image was motion-corrected, and co-registered to a human brain T2-weighted MRI template (MRI space), using PMOD. An already existing brain VOI template was used (AAL-VOIs) for the following regions: caudate, putamen, pallidum, thalamus, and cerebellum. Pineal gland and raphe nuclei were outlined by threshold-based segmentation in the last time frame of each PET scan. All other brain regions contained in the AAL-VOIs atlas were excluded due to very low uptake, determined by preliminary analyses. For pharmacokinetic analyses, an image-derived input function (IDIF) VOI was used, in replacement of blood sampling data. IDIF VOI was generated by threshold-based segmentation of the beginning of the middle cerebral artery, which could be seen in most of the frames corresponding to the first minute after tracer injection.

Pharmacokinetic modelling of dynamic PET images were performed using the calculated TACs extracted from the regional brain VOIs. For the analysis, reversible and irreversible models with IDIF as a surrogate of blood input were evaluated first. For reversible one-tissue (1TCM) and 2-tissue (2TCM) compartmental models, respectively, two (K_1_ and k_2_) and four (K_1_-k_4_) rate constants were determined. The volume of distribution (V_T_) was calculated for the reversible models as V_T_ = (K_1_ /k_2_)(1 + k_3_ /k_4_). Three constants were calculated for the irreversible 2TCM (K_1_-k_3_), and the influx rate (k_i_) was calculated as k_i_ = (K_1_ * k_3_) /(k_2_ + k_3_). The linearized models Logan (reversible) and Patlak (irreversible) graphical analyses were performed with a t* of 25 min and 10 min, respectively. In addition, a simplified reference tissue model analysis was performed using the cerebellum TAC as the input function. The non-displaceable binding potential (BP_nd_) in the SRTM was calculated with the formula BP_nd_ = (k_2_/k_2a_)-1. The Akaike information criterion (AIC), Chi-square (χ^2^), Sum of squares of residuals (SSR) and Coefficient of determination (R^2^) were calculated to determine the fitness of each model.

Additional static PET image analyses were performed with merged images of scans with 60-90 min, and 90-120 min after tracer injection. The percentage of injected dose per cubic centimeter of tissue (%ID/cc) was first calculated for both images and all selected regional brain VOIs. Consequently, standardized uptake volume ratio (SUVr) was obtained for both images, by dividing the %ID/cc of each VOI by the values calculated for the reference region (cerebellum). Average images were generated for both %ID/cc and SUVr images of both timeframes using the Statistical Parametric Mapping software (SPM 25).

### Statistical analyses

Statistical analyses were performed with GraphPad Prism version 10 using standard statistical testing and are represented as mean value ± standard deviation. Blood half-life was calculated using a two-phase decay fit. Linear regression was used to determine the correlation between different models, calculating the Pearson’s coefficient with one-tail. A correlation matrix showing the correlation coefficient was generated. *P*-values < 0.05 were considered statistically significant according to the software (*: *p* < 0.05, **: *p* < 0.01, ***: *p* < 0.001, ****: *p* < 0.0001).

## Results and Discussion

### Biodistribution and dosimetry

The kinetic values of the source organs showed highest mean residence times in the liver (0.183 ± 0.035 h) and cortical bone (0.173 ± 0.041 h) (**Error! Not a valid bookmark self-reference**.). The former points confirm radiotracer metabolism and excretion by hepatobiliary pathways, as the gastrointestinal tract also exhibits moderate residence times. The long residence time in cortical bone points towards uptake of 7-[^18^F]FTrp in the bone, which could be attributed to defluorination of the radiotracer but is more likely to be uptake in the bone marrow, an organ with a high expression of the L-type animo acid transporter (LAT1), the main transporter of Trp.^22^

The organ doses showed the highest mean dose to the pancreas (0.0683 ± 0.019 mSv/MBq), which could be limiting for toxicity (**Error! Not a valid bookmark self-reference**.). As the pancreas is a metabolic active organ which requires Trp as a building block in protein synthesis and also expresses the LAT1^22^, this observation was expected. Similarly, the salivary glands (0.0246 ± 0.008 mSv/MBq) produce a wide range of digestive and other proteins, explaining the high need for Trp. Additionally, the pancreas possesses a local serotonergic system in form of the enterochromaffin cells within the islets of Langerhans, which likely uses 7-[^18^F]FTrp for serotonin synthesis.^23^ Notably, the brain dose is considerably low, which is advantageous for visualizing the serotonergic system in the brain with low background signal from surrounding brain tissue.

The mean effective dose (14.1 ± 0.2 μSv/MBq,**The organ** doses showed the highest mean dose to the pancreas (0.0683 ± 0.019 mSv/MBq), which could be limiting for toxicity (**Error! Not a valid bookmark self-reference**.). As the pancreas is a metabolic active organ which requires Trp as a building block in protein synthesis and also expresses the LAT122, this observation was expected. Similarly, the salivary glands (0.0246 ± 0.008 mSv/MBq) produce a wide range of digestive and other proteins, explaining the high need for Trp. Additionally, the pancreas possesses a local serotonergic system in form of the enterochromaffin cells within the islets of Langerhans, which likely uses 7-[^18F^]FTrp for serotonin synthesis.^23^ Notably, the brain dose is considerably low, which is advantageous for visualizing the serotonergic system in the brain with low background signal from surrounding brain tissue.

**Table *4*)** is at the lower end of the typical range for comparable ^18^F-labeled radiotracers (average effective dose 20.6 ± 6.8 μSv/MBq)^24^, resulting in an effective dose of 3.17 mSv when injecting 225 MBq of 7-[^18^F]FTrp (overall mean in this study).

**Table 3.**
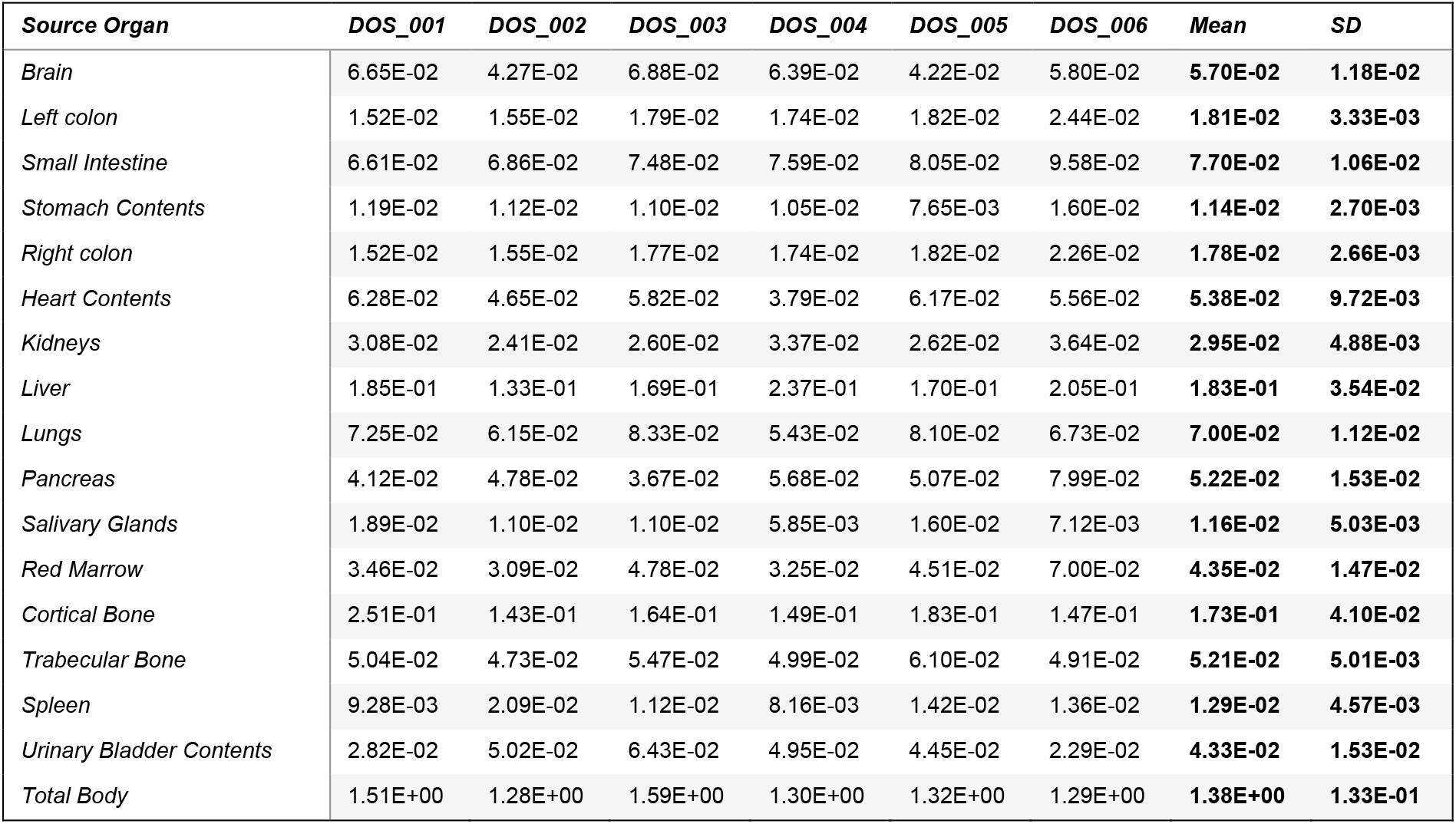
Kinetic values of the source organs calculated by OLINDA (MBq-h/MBq).

**Table 4:** Organ dose and effective dose calculated by OLINDA (mSv/MBq)

| Target Organ | DOS_001 | DOS_002 | DOS_003 | DOS_004 | DOS_005 | DOS_006 | Mean | SD |
| --- | --- | --- | --- | --- | --- | --- | --- | --- |
| Adrenals | 1.63E-02 | 1.13E-02 | 1.59E-02 | 1.55E-02 | 1.51E-02 | 1.70E-02 | 1.52E-02 | 2.17E-03 |
| Brain | 1.32E-02 | 6.98E-03 | 1.22E-02 | 1.06E-02 | 8.93E-03 | 1.01E-02 | 1.03E-02 | 1.95E-03 |
| Breasts | 0.00E+00 | 0.00E+00 | 7.98E-03 | 6.48E-03 | 0.00E+00 | 6.79E-03 | 7.08E-03 | 3.92E-03 |
| Esophagus | 1.22E-02 | 8.24E-03 | 1.20E-02 | 1.03E-02 | 1.12E-02 | 1.12E-02 | 1.09E-02 | 1.44E-03 |
| Eyes | 8.28E-03 | 5.26E-03 | 8.38E-03 | 6.75E-03 | 6.91E-03 | 6.86E-03 | 7.07E-03 | 1.10E-03 |
| Gallbladder Wall | 1.65E-02 | 1.10E-02 | 1.45E-02 | 1.43E-02 | 1.52E-02 | 1.50E-02 | 1.44E-02 | 1.72E-03 |
| Left colon | 2.94E-02 | 2.59E-02 | 2.99E-02 | 2.75E-02 | 3.20E-02 | 3.54E-02 | 3.00E-02 | 3.75E-03 |
| Small Intestine | 2.85E-02 | 2.52E-02 | 3.32E-02 | 3.24E-02 | 3.13E-02 | 3.92E-02 | 3.16E-02 | 5.00E-03 |
| Stomach Wall | 1.81E-02 | 1.41E-02 | 1.69E-02 | 1.54E-02 | 1.60E-02 | 1.92E-02 | 1.66E-02 | 1.90E-03 |
| Right colon | 2.11E-02 | 1.72E-02 | 2.13E-02 | 1.97E-02 | 2.19E-02 | 2.31E-02 | 2.07E-02 | 2.28E-03 |
| Rectum | 1.12E-02 | 8.24E-03 | 1.08E-02 | 1.06E-02 | 1.05E-02 | 9.99E-03 | 1.02E-02 | 1.04E-03 |
| Heart Wall | 2.51E-02 | 1.72E-02 | 2.70E-02 | 1.94E-02 | 2.39E-02 | 2.49E-02 | 2.29E-02 | 4.05E-03 |
| Kidneys | 2.54E-02 | 1.63E-02 | 2.13E-02 | 2.42E-02 | 2.22E-02 | 2.66E-02 | 2.27E-02 | 3.84E-03 |
| Liver | 2.91E-02 | 1.73E-02 | 2.74E-02 | 3.41E-02 | 2.67E-02 | 3.16E-02 | 2.77E-02 | 6.42E-03 |
| Lungs | 1.66E-02 | 1.10E-02 | 1.80E-02 | 1.31E-02 | 1.70E-02 | 1.53E-02 | 1.52E-02 | 2.86E-03 |
| Ovaries | 0.00E+00 | 0.00E+00 | 1.13E-02 | 1.01E-02 | 0.00E+00 | 1.02E-02 | 1.05E-02 | 5.79E-03 |
| Pancreas | 6.18E-02 | 5.37E-02 | 5.18E-02 | 7.13E-02 | 7.24E-02 | 9.86E-02 | 6.83E-02 | 1.89E-02 |
| Prostate | 1.09E-02 | 8.23E-03 | 0.00E+00 | 0.00E+00 | 1.03E-02 | 0.00E+00 | 9.81E-03 | 5.13E-03 |
| Salivary Glands | 4.02E-02 | 1.92E-02 | 2.42E-02 | 1.39E-02 | 3.36E-02 | 1.64E-02 | 2.46E-02 | 7.79E-03 |
| Red Marrow | 1.34E-02 | 8.75E-03 | 1.40E-02 | 1.14E-02 | 1.33E-02 | 1.42E-02 | 1.25E-02 | 2.29E-03 |
| Osteogenic Cells | 2.68E-02 | 1.47E-02 | 1.93E-02 | 1.61E-02 | 2.30E-02 | 1.76E-02 | 1.96E-02 | 3.21E-03 |
| Spleen | 1.81E-02 | 2.39E-02 | 1.97E-02 | 1.56E-02 | 2.30E-02 | 2.24E-02 | 2.05E-02 | 3.36E-03 |
| Testes | 8.21E-03 | 5.71E-03 | 0.00E+00 | 0.00E+00 | 7.27E-03 | 0.00E+00 | 7.06E-03 | 3.60E-03 |
| Thymus | 1.13E-02 | 7.53E-03 | 1.17E-02 | 9.09E-03 | 1.04E-02 | 9.97E-03 | 1.00E-02 | 1.55E-03 |
| Thyroid | 9.65E-03 | 6.44E-03 | 9.18E-03 | 7.24E-03 | 8.57E-03 | 7.61E-03 | 8.12E-03 | 1.08E-03 |
| Urinary Bladder Wall | 2.28E-02 | 2.92E-02 | 8.79E-03 | 3.33E-02 | 2.95E-02 | 1.94E-02 | 2.38E-02 | 9.95E-03 |
| Uterus | 0.00E+00 | 0.00E+00 | 1.13E-02 | 1.13E-02 | 0.00E+00 | 1.07E-02 | 1.11E-02 | 6.08E-03 |
| Total Body | 1.08E-02 | 7.17E-03 | 1.11E-02 | 9.72E-03 | 9.64E-03 | 1.02E-02 | 9.77E-03 | 1.46E-03 |
| Effective Dose (ICRP-103 ) | 1.45E-02 | 1.10E-02 | 1.47E-02 | 1.42E-02 | 1.44E-02 | 1.58E-02 | 1.41E-02 | 1.62E-03 |

In the PET images, 7-[^18^F]FTrp showed rapid distribution through circulation following administration (**Figure 1A, C, Suppl. Figure 1**). Accumulation in the bladder, liver and kidney confirmed mainly hepatobiliary clearance. However, liver accumulation could be partially attributed to ongoing kynurenine metabolism, since the enzyme tryptophan 2,3 deoxygenase (TDO), the first and rate-limiting step in tryptophane metabolism, is mainly expressed in the liver and responsible for the majority of Trp metabolism.^25^ Notably, strong accumulation of 7-[^18^F]FTrp in the pancreas was observed as already indicated in the dosimetry analysis (**Figure 1B, Suppl. Figure 2**). The blood-half-life was calculated in GraphPad prism using the TACs and confirmed rapid distribution (half-life fast: 256 s ± 44 s, half-life slow: 1362 s ± 153, n = 5, 1 data set was excluded due to the software not being able to calculate a confidence interval).

**Figure 1:**
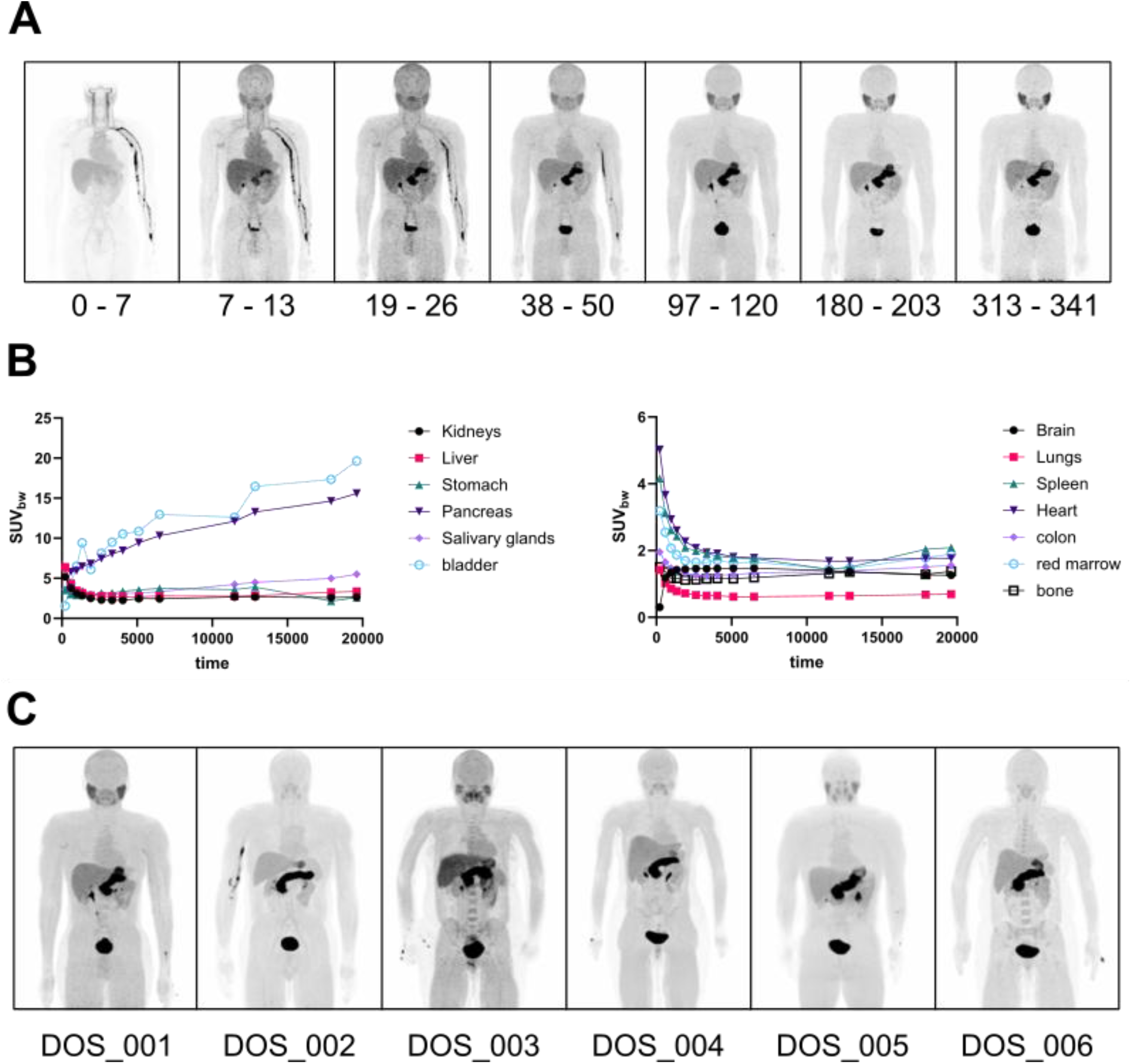
Biodistribution of 7-[^18^F]FTrp in healthy volunteers. **A** Representative maximum intensity projection (MIP) of one healthy volunteer at different time points after injection. **B** TACs of source organs separated in organs with higher activity (left) and lower activity (right). **C** MIPs of all six healthy volunteers at 75 – 120 min post-injection.

### PET brain dynamic analysis

In the brain, 7-[^18^F]FTrp presented with visual accumulation in the pineal gland in the PET images, the main areas of serotonin production in the brain, and in the TACs (**Figure 2A, Suppl. Figure 3**). Pharmacokinetic analysis in different models confirmed regional brain uptake of 7-[^18^F]FTrp. In all models, except SRTM, all brain areas showed a higher uptake (V_T_, or K_i_, but not BP_nd_) compared to the cerebellum, which had the lowest uptake, and was thus used as a reference region (**Table 5**).

**Figure 2:**
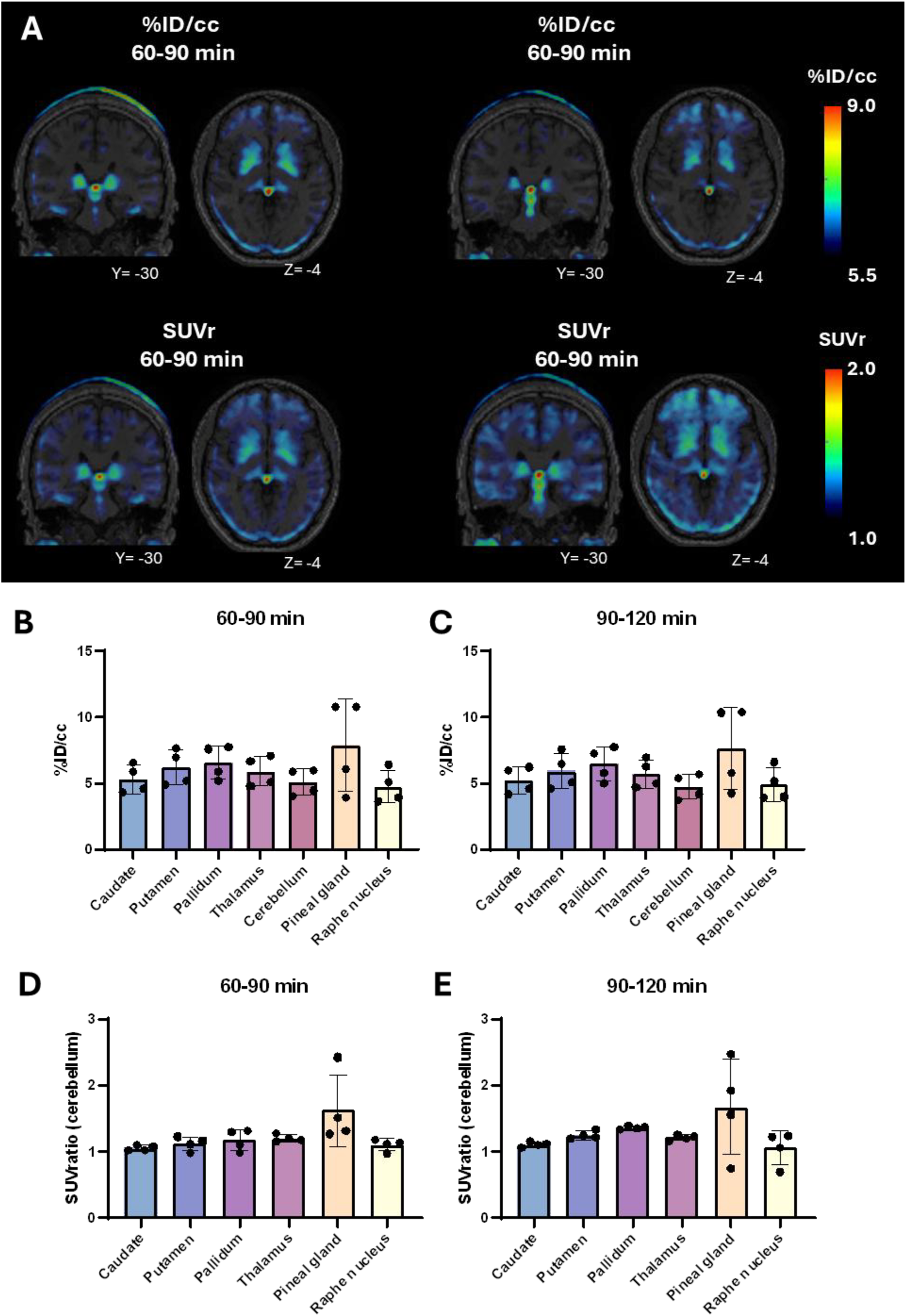
7-[^18^F]FTrp brain regional uptake in static analyses. **A**. Merged images of radiotracer uptake. **B-D**. Graphical representation of brain uptake.

**Table 5:** Outcome parameters calculated for each of the kinetic models, in the selected brain regions.

| *Brain region* | *Reversible*<br>*1TCM ($V_T$)* | *Reversible*<br>*2TCM ($V_T$)* | *Irreversible*<br>*2TCM ($K_i$)* | *Logan*<br>*($V_T$)* | *Patlak*<br>*($K_i$)* | *SRTMcer*<br>*($BP_{ND}$)* |
| --- | --- | --- | --- | --- | --- | --- |
| *Caudate* | 1.12 ± 0.12 | 1.16 ± 0.11 | 0.0034 ± 0.0001 | 1.16 ± 0.11 | 0.0036 ± 0.0002 | 0.05 ± 0.04 |
| *Putamen* | 1.28 ± 0.17 | 1.33 ± 0.17 | 0.0038 ± 0.0013 | 1.33 ± 0.17 | 0.0039 ± 0.0014 | 0.16 ± 0.01 |
| *Pallidum* | 1.40 ± 0.14 | 1.45 ± 0.11 | 0.0045 ± 0.0012 | 1.44 ± 0.13 | 0.0048 ± 0.0014 | 0.25 ± 0.06 |
| *Pineal gland* | 1.01 ± 0.11 | 1.08 ± 0.10 | 0.0034 ± 0.0004 | 1.09 ± 0.10 | 0.0035 ± 0.0005 | 0.25 ± 0.50 |
| *Raphe*<br>*nucleus* | 1.62 ± 0.81 | 2.52 ± 0.90 | 0.0090 ± 0.0043 | 1.74 ± 0.72 | 0.0096 ± 0.0049 | 0.80 ± 0.70 |
| *Cerebellum* | 0.99 ± 0.27 | 1.12 ± 0.27 | 0.0018 ± 0.0008 | 1.07 ± 0.21 | 0.0018 ± 0.0008 | 0.10 ± 0.09 |

To determine the best fitting model, the goodness of fit outcome parameters AIC, χ^2^, SSR and R^2^ were compared between models. For all outcome parameters, except R^2^, Patlak showed the most suitable numbers (Suppl. Table 1). These data confirmed the irreversible nature of 7-[^18^F]FTrp and suggested the use of Patlak as the main kinetic modelling PET imaging analyses with this tracer.

### PET brain static uptake and correlation analyses

Static PET imaging regional brain analyses corresponded to dynamic results, showing uptake of 7-[^18^F]FTrp in all aforementioned areas. This uptake was similar between %ID/cc calculation of 60-90 min with 90-120 min, and between SUVr calculations of 60-90 min with 90-120 min (**Table 6, Figure 2B**).

**Table 6:** 7-[^18^F]FTrp brain regional uptake in static analyses.

| *Brain region* | *%ID/cc (60–90 min)* | *%ID/cc (90–120 min)* | *SUVr (cer) 60–90 min* | *SUVr (cer) 90–120 min* |
| --- | --- | --- | --- | --- |
| *Caudate* | 5.34 ± 1.08 | 5.24 ± 1.02 | 1.05 ± 0.04 | 1.11 ± 0.04 |
| *Putamen* | 6.19 ± 1.34 | 5.95 ± 1.36 | 1.12 ± 0.10 | 1.25 ± 0.06 |
| *Pallidum* | 6.61 ± 1.28 | 6.46 ± 1.26 | 1.18 ± 0.16 | 1.36 ± 0.02 |
| *Pineal gland* | 7.91 ± 3.48 | 7.71 ± 3.15 | 1.63 ± 0.54 | 1.68 ± 0.72 |
| *Raphe nucleus* | 4.77 ± 1.26 | 4.92 ± 1.27 | 1.10 ± 0.09 | 1.05 ± 0.25 |
| *Cerebellum* | 5.10 ± 1.02 | 4.75 ± 0.94 | 1.00 ± 0.00 | 1.00 ± 0.00 |

To determine if the SRTM model, which does not require arterial input, and if the static models can represent the results obtained from the Patlak analyses, a correlation matrix was performed (**Figure 3**). Results showed that both SRTM and SUVr 60-90 min models had the highest correlation coefficients with Patlak, suggesting the feasibility of using simplified scan protocols and analytical methods.

**Figure 3:**
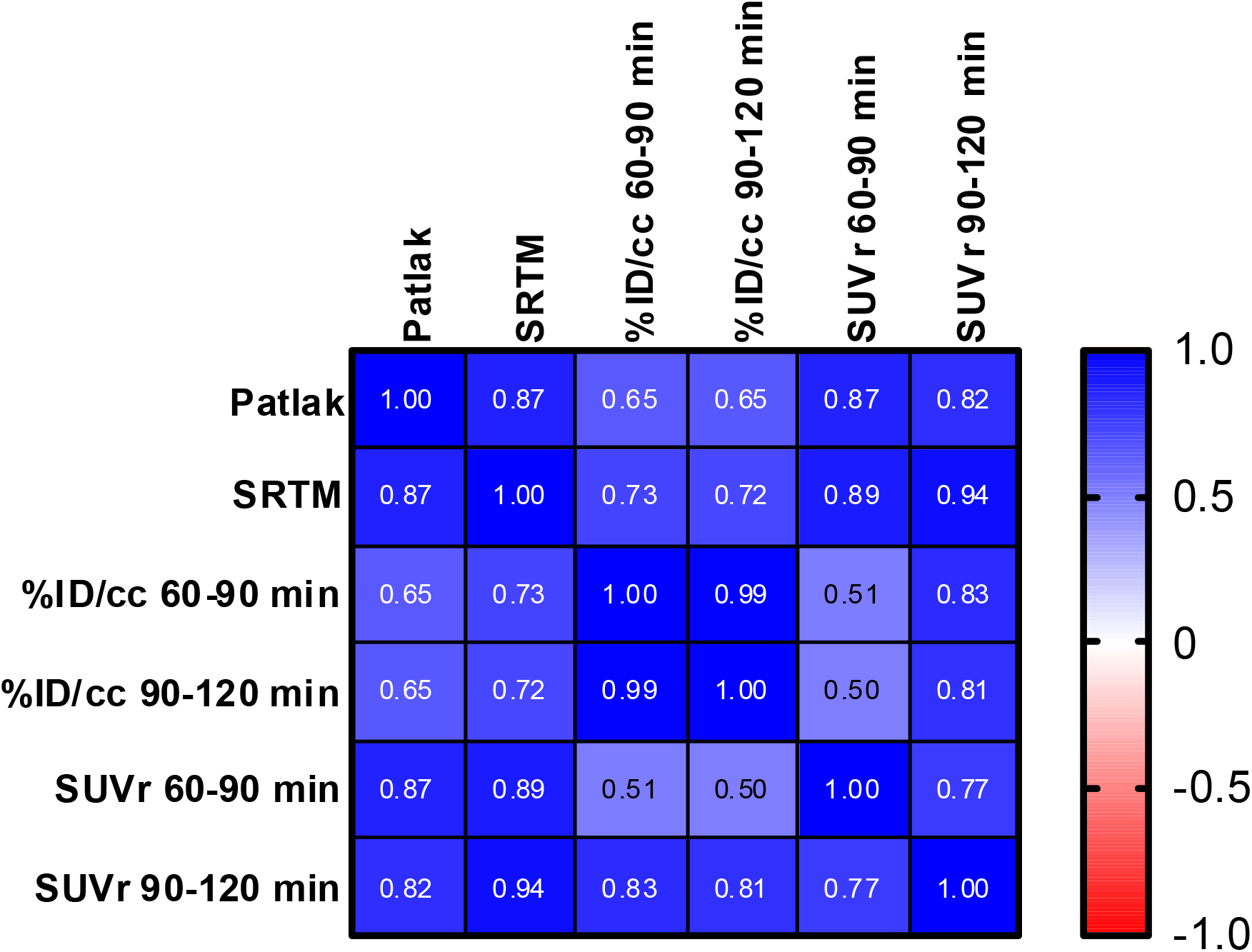
Correlation matrix of different dynamic and static PET imaging analytical models.

## Conclusion

This proof-of-principle study investigated the biodistribution and dosimetry profile of 7-[^18^F]FTrp in humans and its ability to serve as a marker for serotonin synthesis. Overall, the dosimetry showed a favorable safety profile within the range of comparable ^18^F-labeled radiotracers. Notable uptake of 7-[^18^F]FTrp was observed in excreting organs like liver and bladder, the gastrointestinal tract and pancreas, salivary glands and bone. In consequence, pancreas and salivary glands are likely the dose-limiting organs in further studies. These observations are explainable by the various metabolic pathways in which Trp is involved to undergo biosynthesis of kynurenine or serotonin. The involvement of altered amino acid metabolism in diseases like cancer or Parkinson’s disease warrants the additional investigation of 7-[^18^F]FTrp in this context.^17^

The dynamic brain study showed promising accumulation of 7-[^18^F]FTrp in the pineal gland and the raphe nuclei at later time points, pointing towards the capability of the radiotracer to visualize serotonin synthesis from Trp. The observed inter-subject variability is encouraging to explore 7-[^18^F]FTrp in a more sophisticated study, e.g. comparing patients with altered serotonin metabolism with healthy controls. Ultimately, 7-[18F]FTrp may offer a powerful means to explore serotonin dysfunction in vivo, aiding in the diagnosis and management of diseases involving serotoninergic dysregulation.

