## Supplemental Information for "Preliminary First-in-Human Pharmacokinetic Evaluation, Dosimetry and Safety of 7-[^18^F]-Fluorotryptophan as PET Imaging Agent to Visualize Serotonin Synthesis in the Brain"

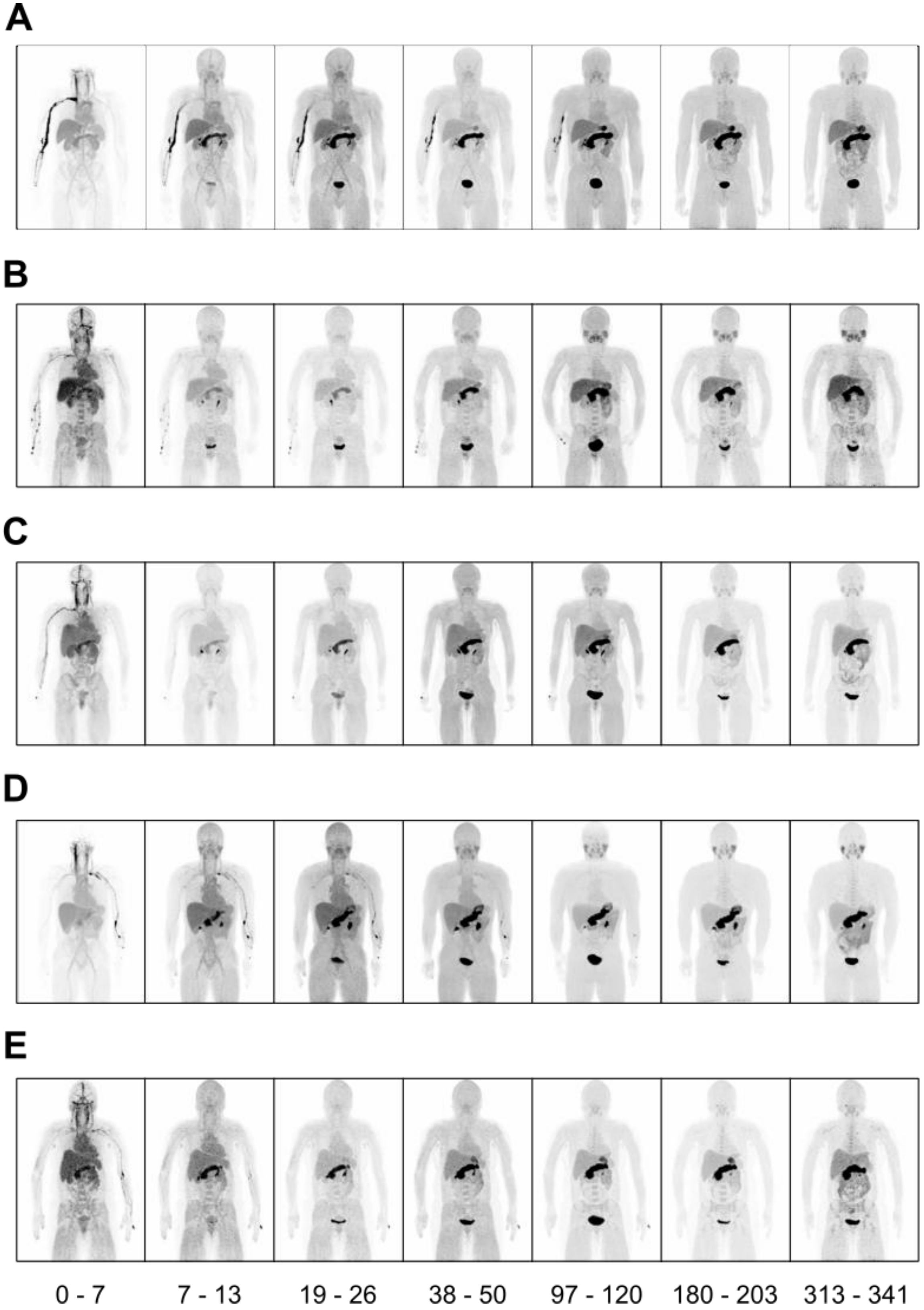


**Supplemental Figure 1: Biodistribution of 7-[$^{18}$F]FTrp in healthy volunteers. A-E** Representative maximum intensity projections (MIPs) of healthy volunteers DOS_002 – DOS_006 at different time points after injection.

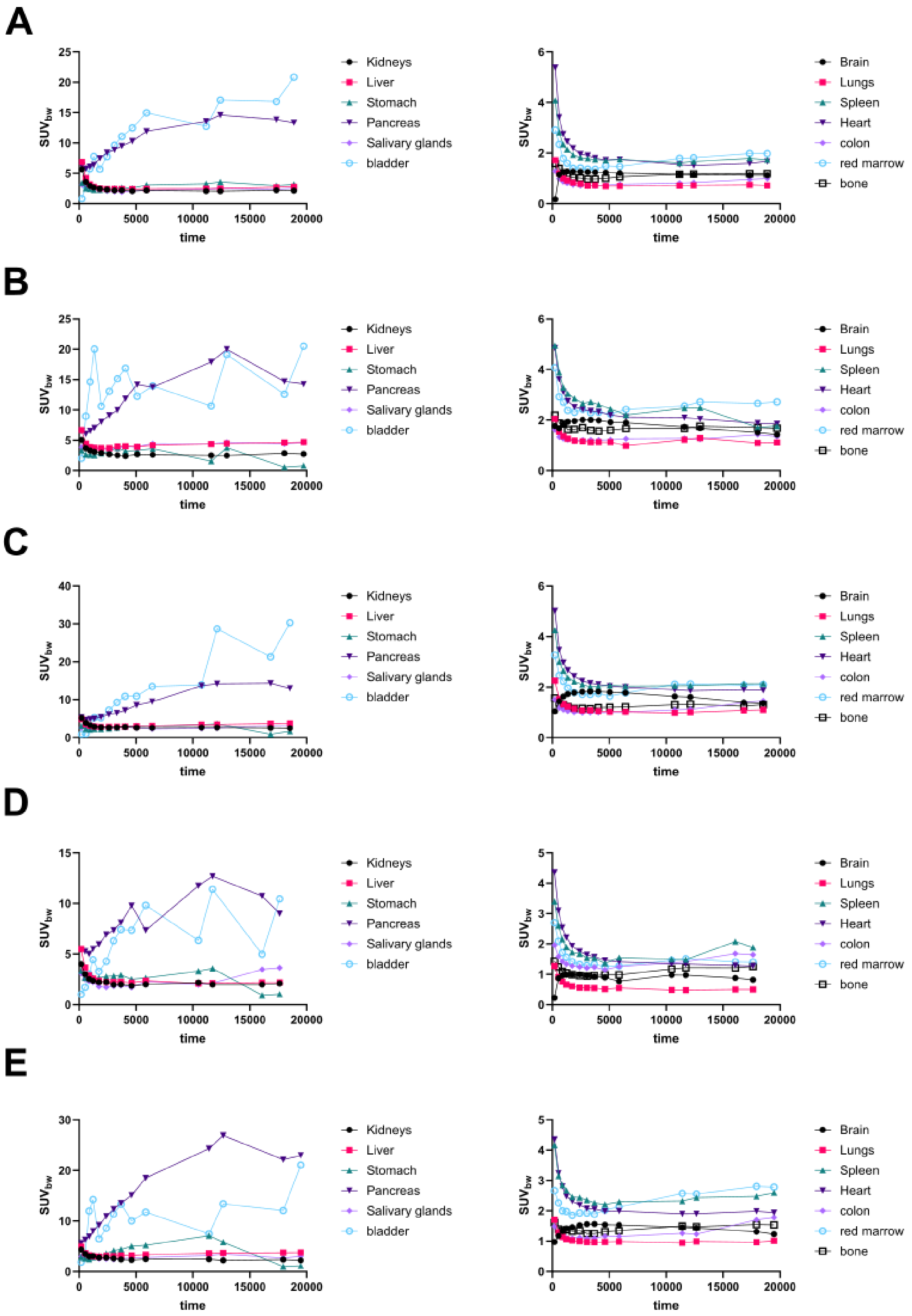


**Supplemental Figure 2: TACs of source organs. A – E** DOS_002 – DOS_006 separated in organs with higher activity (left) and lower activity (right).

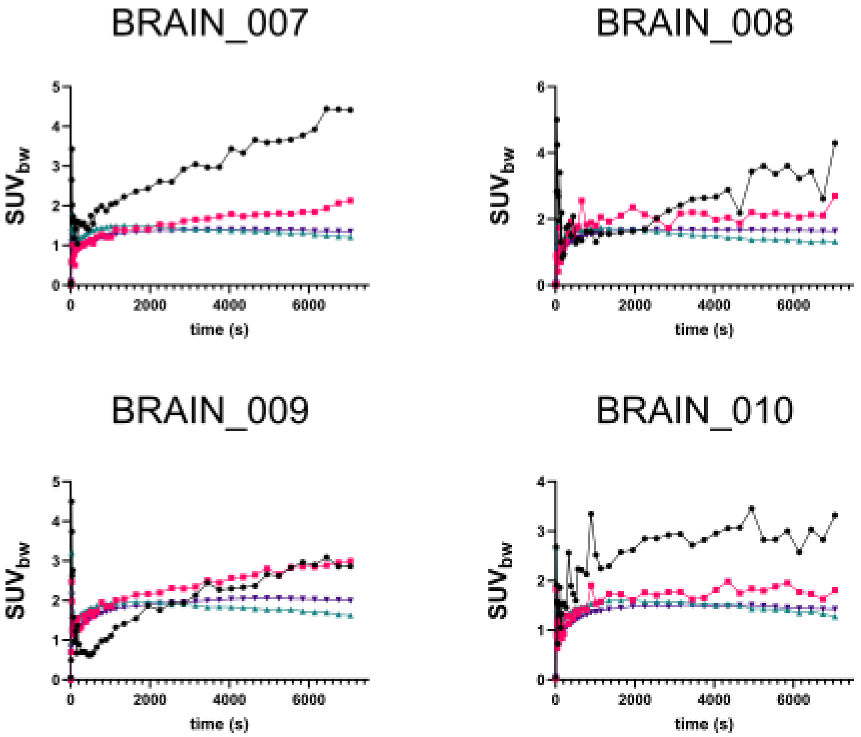


**Supplemental Figure 3: TACs of the dynamic brain PET scans.**

**Supplemental Table 1: Goodness of fit parameters comparison between dynamic models.**

| | ***Caudate*** | ***Putamen*** | ***Pallidum*** | ***Pineal gland*** | ***Raphe nucleus*** | ***Cerebellum*** |
|---|---|---|---|---|---|---|
| ***AIC*** | | | | | | |
| *Reversible 1TCM (Vt)* | 92.6 ± 23.1 | 92.5 ± 10.7 | 80.0 ± 6.6 | 162.4 ± 37.7 | 163.3 ± 26.3 | 149.1 ± 8.9 |
| *Reversible 2TCM (Vt)* | 50.3 ± 51.7 | 57.5 ± 32.9 | 40.5 ± 34.7 | 141.9 ± 55.3 | 150.5 ± 34.8 | 83.5 ± 51.5 |
| *Irreversible 2TCM (Ki)* | 83.8 ± 28.5 | 81.7 ± 17.7 | 66.5 ± 12.6 | 161.5 ± 46.1 | 160.6 ± 29.4 | 145.0 ± 11.6 |
| *Logan (Vt)* | 149.7 ± 3.7 | 141.8 ± 5.1 | 158.9 ± 15.6 | 190.2 ± 3.9 | 124.5 ± 12.6 | 121.1 ± 4.0 |
| ***Patlak (Ki)*** | **-116.4 ± 9.5** | **-108.5 ± 6.0** | **-101.6 ± 9.5** | **-82.0 ± 44.1** | **-127.7 ± 29.8** | **-124.0 ± 10.2** |
| *SRTMcer (BP_ND)* | 9.1 ± 18.2 | -8.6 ± 25.4 | 18.5 ± 44.7 | 93.1 ± 36.4 | 112.4 ± 55.0 | 97.7 ± 126.7 |
| ***$X^2$*** | | | | | | |
| *Reversible 1TCM (Vt)* | 8.81 ± 5.41 | 8.02 ± 2.13 | 4.41 ± 2.79 | 53.18 ± 55.82 | 44.42 ± 24.56 | 28.77 ± 6.16 |
| *Reversible 2TCM (Vt)* | 5.18 ± 7.14 | 4.20 ± 3.71 | 2.91 ± 2.63 | 42.99 ± 58.71 | 34.53 ± 26.34 | 10.42 ± 13.23 |
| *Irreversible 2TCM (Ki)* | 7.53 ± 5.81 | 6.41 ± 2.99 | 4.39 ± 1.39 | 60.26 ± 76.45 | 42.27 ± 26.13 | 25.97 ± 7.45 |
| *Logan (Vt)* | 2334.6 ± 496.7 | 1552.5 ± 380.7 | 4598.9 ± 2876.8 | 19728.4 ± 4090.3 | 740.6 ± 604.7 | 519.1 ± 106.8 |
| ***Patlak (Ki)*** | **0.003 ± 0.002** | **0.005 ± 0.003** | **0.007 ± 0.004** | **0.040 ± 0.034** | **0.003 ± 0.003** | **0.002 ± 0.001** |
| *SRTMcer (BP_ND)* | 1.23 ± 0.47 | 0.87 ± 0.46 | 1.90 ± 1.28 | 10.10 ± 8.67 | 19.29 ± 17.71 | 57.49 ± 90.52 |
| ***SSR*** | | | | | | |
| *Reversible 1TCM (Vt)* | 18.32 ± 18.8 | 21.92 ± 12.5 | 15.4 ± 6.8 | 196.0 ± 155.2 | 354.9 ± 352.1 | 79.6 ± 39.4 |
| *Reversible 2TCM (Vt)* | 12.12 ± 9.8 | 12.02 ± 13.5 | 8.52 ± 10.0 | 146.2 ± 164.1 | 284.2 ± 339.3 | 32.3 ± 44.8 |
| *Irreversible 2TCM (Ki)* | 16.0 ± 18.6 | 17.6 ± 13.2 | 11.7 ± 7.6 | 209.2 ± 213.6 | 337.8 ± 361.0 | 71.2 ± 40.1 |
| *Logan (Vt)* | 39688.6 ± 8444.3 | 26392.9 ± 6472.1 | 78181.0 ± 48906.0 | 335383.6 ± 69536.7 | 12590.6 ± 10279.6 | 8825.5 ± 1815.2 |
| ***Patlak (Ki)*** | **0.06 ± 0.05** | **0.09 ± 0.07** | **0.13 ± 0.10** | **0.79 ± 0.73** | **0.05 ± 0.06** | **0.04 ± 0.03** |
| *SRTMcer (BP_ND)* | 2.18 ± 1.33 | 1.94 ± 0.60 | 4.79 ± 4.50 | 35.93 ± 22.81 | 127.55 ± 123.34 | 118.57 ± 159.57 |
| ***$R^2$*** | | | | | | |
| *Reversible 1TCM (Vt)* | 0.78 ± 0.18 | 0.77 ± 0.11 | 0.88 ± 0.03 | 0.50 ± 0.19 | 0.67 ± 0.21 | -0.30 ± 0.42 |
| *Reversible 2TCM (Vt)* | 0.87 ± 0.20 | 0.87 ± 0.13 | 0.94 ± 0.06 | 0.69 ± 0.15 | 0.75 ± 0.21 | 0.52 ± 0.66 |
| *Irreversible 2TCM (Ki)* | 0.82 ± 0.18 | 0.81 ± 0.13 | 0.92 ± 0.04 | 0.53 ± 0.15 | 0.70 ± 0.22 | -0.16 ± 0.46 |
| *Logan (Vt)* | 0.99 ± 0.00 | 0.99 ± 0.00 | 0.99 ± 0.00 | 0.99 ± 0.00 | 0.99 ± 0.00 | 0.99 ± 0.00 |
| ***Patlak (Ki)*** | **0.76 ± 0.12** | **0.60 ± 0.27** | **0.70 ± 0.22** | **0.73 ± 0.19** | **0.40 ± 0.25** | **0.13 ± 0.18** |
| *SRTMcer (BP_ND)* | 0.97 ± 0.01 | 0.98 ± 0.01 | 0.96 ± 0.03 | 0.87 ± 0.08 | 0.87 ± 0.10 | -1.27 ± 3.36 |
